# Sex differences in pain-related behaviors and clinical progression of disease in mouse models of visceral pain

**DOI:** 10.1101/2022.01.11.475721

**Authors:** Adela M. Francis-Malave, Santiago Martinez Gonzalez, Caren Pichardo, Torri D. Wilson, Luis G. Rivera, Lauren R. Brinster, Yarimar Carrasquillo

**Affiliations:** National Center for Complementary and Integrative Health, National Institutes of Health, Bethesda, MD, United States; Office of Research Services, Division of Veterinary Resources, National Institutes of Health, Bethesda, MD, United States

**Keywords:** visceral pain, sex differences, colitis, dextran sulfate sodium (DSS), intracolonic capsaicin, referred abdominal hypersensitivity

## Abstract

Previous studies have reported sex differences in irritable bowel syndrome (IBS) and inflammatory bowel disease (IBD) patients, including differences in visceral pain perception. Despite this, sex differences in behavioral manifestations of visceral pain and underlying pathology of the gastrointestinal tract have been largely understudied in preclinical research. In this study, we evaluated potential sex differences in spontaneous visceral nociceptive responses, referred abdominal hypersensitivity, disease progression and bowel pathology in mouse models of acute and persistent colon inflammation. Our experiments show that females exhibit more visceral nociceptive responses and referred abdominal hypersensitivity than males in the context of acute but not persistent colon inflammation. We further demonstrate that, following acute and persistent colon inflammation, visceral pain-related behavioral responses in females and males are distinct, with increases in licking of the abdomen only observed in females and increases in abdominal contractions only seen in males. During persistent colon inflammation, males exhibit worse disease progression than females, which is manifested as worse physical appearance and higher weight loss. However, no measurable sex differences were observed in persistent inflammation-induced bowel pathology, stool consistency or fecal blood. Overall, our findings demonstrate that visceral pain-related behaviors and disease progression in the context of acute and persistent colon inflammation are sex-dependent, highlighting the importance of considering sex as a biological variable in future mechanistic studies of visceral pain as well as in the development of diagnostics and therapeutic options for chronic gastrointestinal diseases.

## INTRODUCTION

Patients with irritable bowel syndrome (IBS), inflammatory bowel disease (IBD), and other chronic gastrointestinal diseases often manifest altered visceral pain perception that has been related to the development of visceral hypersensitivity [9,34,47]. Importantly, clinical studies have shown that women with IBS exhibit higher sensitivity to repetitive rectal distention and report more severe abdominal pain or discomfort than men, suggesting sex differences in visceral pain perception in IBS [3,4,44]. Sex differences have also been reported in the clinical manifestation, disease course, complications, psychiatric comorbidities, central processing of visceral pain and pathophysiology of IBS and other chronic gastrointestinal disorders [13,26,37,43]. In contrast, a recent study found no sex differences in visceral pain thresholds to rectal balloon distensions in healthy young women and men [19], suggesting that baseline visceral sensitivity is not dependent on sex. Evaluation of the mechanisms underlying sex differences in visceral sensitivity in human subjects is limited by methodological and experimental challenges and are, thus, not completely understood [1,11,16]. Preclinical studies in rodents offer a valuable alternative to evaluate sex-specific factors in chronic gastrointestinal diseases in a more controlled setting, providing insights towards the development of more effective diagnostic and treatment options for patients.

Preclinical studies evaluating sex differences in visceral sensitivity in rodents have predominantly measured visceromotor responses to colorectal distension in rats. These studies have shown that visceral sensitivity in rodents is sex-dependent, with females exhibiting higher visceromotor responses to colonic distension than males at baseline and after acute inflammation of the colon [23, 45]. Sex differences in visceromotor responses to colonic distention have also been reported in the context of stress, where female rats exhibit a stronger relationship between early life adversity and the development of stress-induced visceral hypersensitivity [41]. Separate studies further show that sex hormones contribute to sex differences in stress-induced visceral hypersensitivity, with estradiol facilitating and testosterone decreasing stress effects on visceral responses [21]. Despite this knowledge, sex differences in visceral sensitivity after acute and persistent inflammation of the bowel remains poorly understood. Evaluating and characterizing potential sex differences in visceral pain-related responses in mice under pathological conditions is essential for the evaluation of mechanisms driving sex differences in visceral pain.

In the present study, we systematically evaluated potential sex differences in visceral pain-related responses using two well-characterized mouse models of chemically induced visceral hypersensitivity: intracolonic capsaicin, which elicits transient neurogenic inflammation [28], spontaneous nociceptive responses and referred abdominal hyperalgesia [29, 42]; and the Dextran Sodium Sulfate (DSS) model of colitis, which elicits prolonged inflammation in the bowel as well as visceral and referred abdominal hypersensitivity [2, 10] (**Figure 1**). Parallel experiments evaluated potential sex differences in disease progression of DSS-induced colitis and associated bowel pathology. Our experiments revealed sex differences in pain-related behaviors after acute colonic irritation as well as in the clinical progression of colitis. Visceral pain-related behaviors and colon pathology after persistent inflammation of the bowel, however, were similar in males and females. Collectively, these findings set a foundation for future mechanistic studies of sex differences in visceral pain.

**Figure 1.**
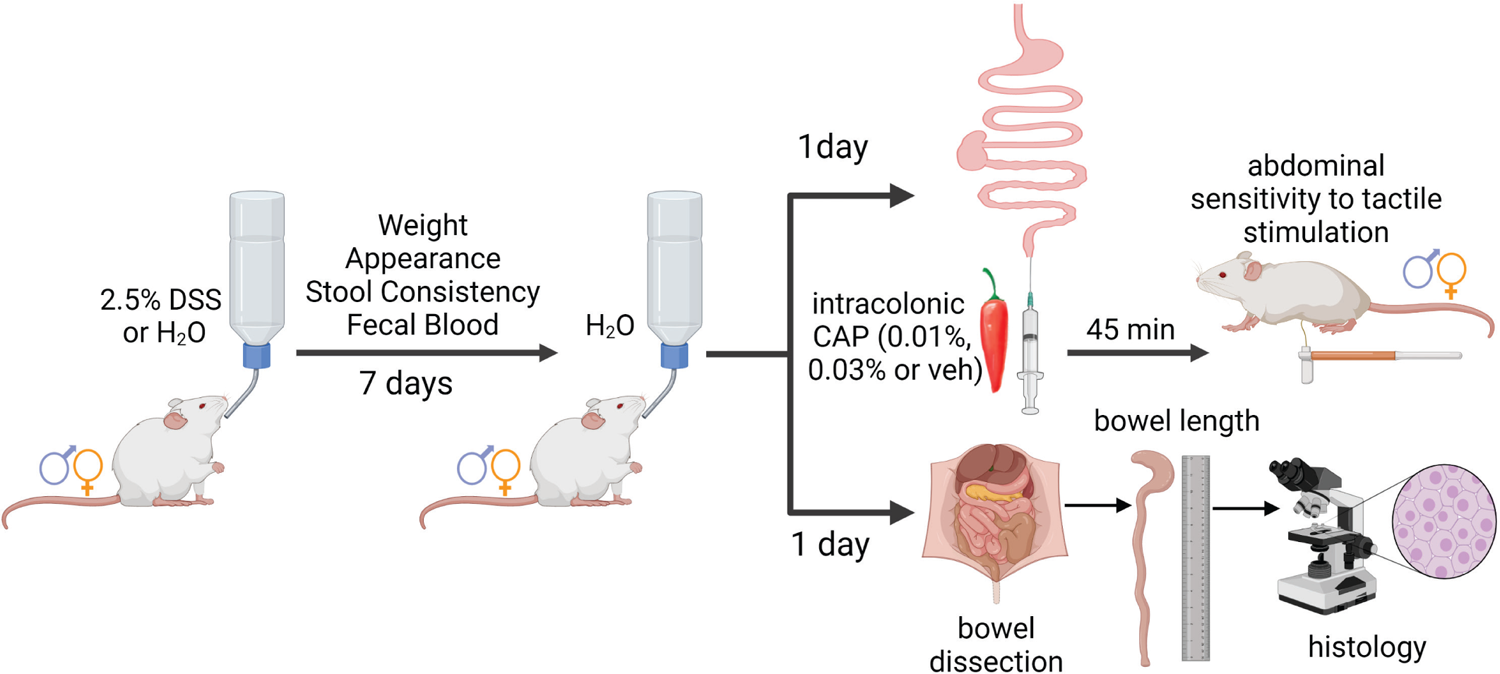
Experimental timeline for behavioral and histological experiments in a model of DSS-induced colitis. Male and female mice were treated with either 2.5% DSS or water and clinical progression of disease was monitored for 7 days. Spontaneous responses to intracolonic capsaicin (0.01% or 0.03%) or vehicle control and referred abdominal sensitivity to tactile stimulation were measured 1d after the end of DSS treatment. Bowels were dissected and measured and histologically processed for pathology assessment.

## RESULTS

### Female mice display more capsaicin-induced spontaneous pain-related behaviors than male mice

To begin evaluating potential sex differences in visceral pain-related behaviors, we used the intracolonic capsaicin model and measured spontaneous nociceptive responses in both male and female mice (**Figure 2A**). Spontaneous capsaicin-induced nociceptive behaviors were defined as licking, stretching, and dragging of the abdomen as well as abdominal contractions. Consistent with previous reports [28, 29], intracolonic application of capsaicin (0.01%) elicited robust pain-related behaviors that were significantly (p < 0.0001) higher than those observed following an intracolonic injection of vehicle in female mice (**Figure 2B**). In marked contrast, however, spontaneous pain-related behaviors were indistinguishable in male mice following the intracolonic administration of 0.01% capsaicin or vehicle control (**Figure 2B**). To determine if male mice can display capsaicin-induced hypersensitivity, we next injected mice with a higher dose of capsaicin (0.03%). As illustrated in **Figure 2C**, intracolonic administration of 0.03% capsaicin elicited significant (p = 0.0010) increases in spontaneous pain-related behaviors in males, compared to vehicle-injected mice. These results demonstrate that capsaicin-induced spontaneous nociceptive responses are sex-dependent, with higher doses of capsaicin required in males than in females to elicit comparable nociceptive behavioral responses.

**Figure 2.**
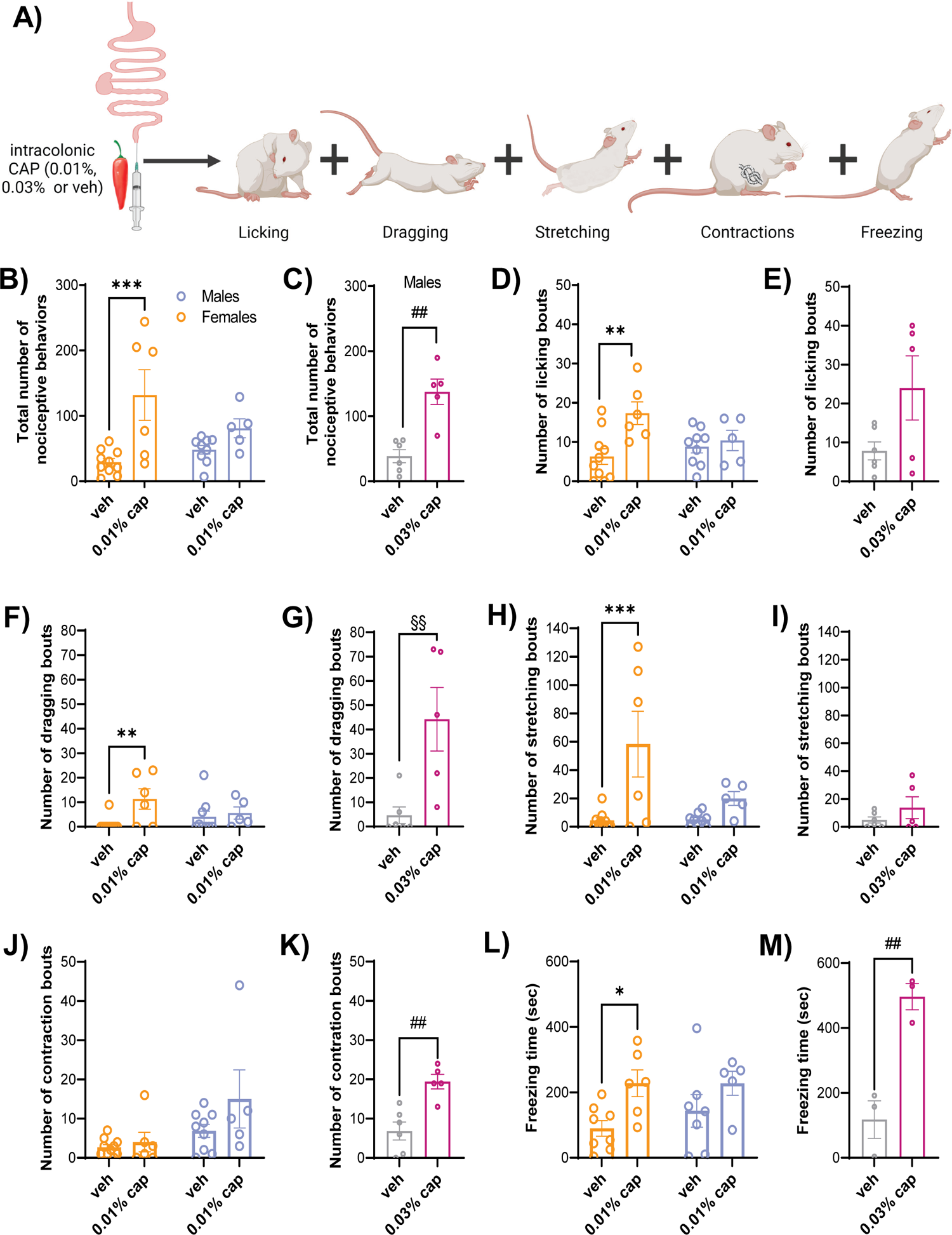
Capsaicin-induced nociceptive behaviors are higher in female than male mice and the behavioral manifestation is sex-dependent. (A) Timeline of behavioral experiments. Male and female mice were injected with intracolonic capsaicin (0.01% or 0.03%) and capsaicin-evoked nociceptive responses, defined as licking, stretching, dragging, contractions of the abdomen, and freezing behaviors were recorded. (B-M) Total number of pain-related behaviors (B-C), licking bouts (D-E), dragging bouts (F-G), stretching bouts (H-I), contraction bouts (J-K) and time spent in freezing behavior (L-M) in vehicle control, 0.01% capsaicin and 0.03% capsaicin. Data is presented as mean ± SEM. n = 6-10 females and 5-9 males. *p < 0.05, **p < 0.01, ***p < 0.001: veh vs 0.01% cap; two-way ANOVA followed by Šídák’s multiple comparisons test; ##p < 0.01: veh vs 0.03% cap; unpaired t-test; §§p < 0.01: veh vs 0.03% cap; Mann Whitney test.

The individual components contributing to the total number of capsaicin-induced nociceptive behaviors were analyzed individually to identify which specific behaviors contribute to the observed sex differences (**Figure 2D-K**). Consistent with the results observed in the cumulative analysis, females injected with 0.01% capsaicin displayed significant increases in most behavioral parameters when compared to control vehicle-injected females. Thus, female mice injected with 0.01% capsaicin displayed a significant (p < 0.01) increase in the number of licking bouts (**Figure 2D**), dragging bouts (**Figure 2F**) and stretching bouts (**Figure 2H**), compared to vehicle-injected females. The number of contraction bouts was the only parameter in females that was indistinguishable between capsaicin- and vehicle-injected mice (**Figure 2J**). In marked contrast and consistent with the results observed in the cumulative behavioral analysis, all the parameters measured were indistinguishable between vehicle- and capsaicin-injected male mice (**Figure 2D, 2F, 2H, 2J**), confirming the lack of measurable behavioral hypersensitivity in male mice following intracolonic injection of 0.01% capsaicin.

Evaluation of the individual behavioral components in male mice injected with a higher dose of capsaicin (0.03%) revealed that the capsaicin-induced increases in behavioral responses observed at this concentration were mainly driven by significant (p < 0.01) increases in abdominal dragging and contraction bouts (**Figure 2G, 2K**) but not by changes in licking or stretching of the abdomen (**Figure 2E, 2I**). Together, these results highlight that the behavioral manifestation of capsaicin-induced visceral hypersensitivity is sexually dimorphic, with increases in licking and stretching of the abdomen observed in females only and increases in abdominal contractions seen solely in male mice. The common behavioral manifestation observed between sexes following intracolonic capsaicin administration was increases in dragging of the abdomen.

Previous studies have indicated that intracolonic capsaicin elicits freezing behaviors in a dose-dependent manner [6]. Consistent with these findings, the time spent freezing was significantly (p < 0.05) higher in females following intracolonic injection of 0.01% capsaicin than in females after the injection of vehicle control (**Figure 2L**). In line with the sex differences presented above, no measurable differences were observed in the time spent freezing in male mice injected with 0.01% capsaicin when compared to vehicle control (**Figure 2L**). Time spent freezing, however, was significantly (p < 0.01) higher in males when a higher dose of capsaicin (0.03%) was used, compared with vehicle control (**Figure 2M**). These results demonstrate that while both males and females exhibit capsaicin-induced freezing, males require a higher dose of capsaicin than females to display freezing behaviors, further confirming that the manifestation of capsaicin-induced nociceptive responses is sex-dependent and that females are more hypersensitive.

### Clinical progression of disease in a DSS-induced mouse model of colitis is sexually dimorphic

The DSS model of colitis was used to evaluate sex differences in clinical progression of disease as well as visceral pain-related responses and referred abdominal sensitivity in mice. Male and female mice were treated with either 2.5% DSS or water, ad libitum for seven days (**Figure 3A**). Disease Activity Index (DAI), defined as a cumulative score based on the percentage of weight loss, stool consistency, presence of fecal blood and physical appearance (**Table 1**), was calculated daily to monitor the clinical progression of colitis. As illustrated in **Figure 3B**, the DAI score was dependent on the dose of DSS administered, with increasing percent of DSS resulting in higher DAI score. The rest of the experiments in this study were performed using 2.5% DSS which consistently elicited disease in all mice but was not at ceiling level.

**Figure 3.**
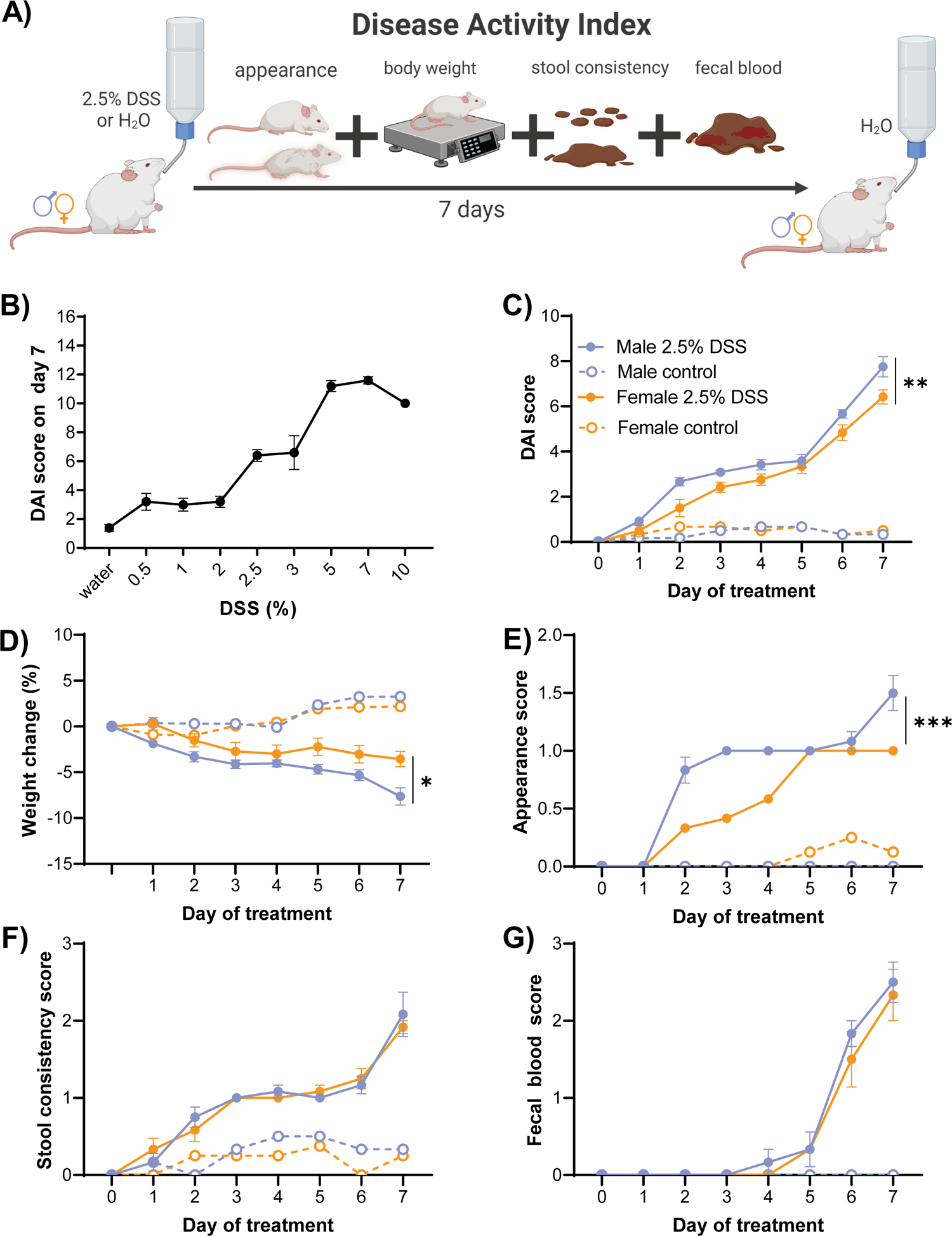
Clinical progression of disease in a DSS-induced mouse model of colitis is sexually dimorphic. (A) Timeline for the induction of experimental colitis. Male and female mice were treated with either 2.5% DSS or water, ad libitum for seven days. Disease Activity Index (DAI) cumulative score was obtained daily to monitor the progression of DSS-induced colitis. (B) DAI score was dependent on the dose of DSS administered. (C-G) Cumulative DAI scores (C) and its individual components, defined as percentage of weight change (D), appearance (E), stool consistency (F) and fecal blood (G) were analyzed. Data is presented as mean ± SEM. n = 6-12 females and 6-12 males. *p < 0.05, **p < 0.01, ***p < 0.001: male vs female; two-way ANOVA.

**Table 1.** Disease Activity Index (DAI). DAI score is the sum of the following components: the percentage of weight loss, stool consistency, fecal blood and appearance. Each individual component was scored according to the descriptions in the table.

| Score | Weight Loss (%) | Stool Consistency | Fecal Blood | Appearance |
| --- | --- | --- | --- | --- |
| 0 | None or gain | Formed and hard | Absence | “Normal” |
| 1 | 1-5 | Formed and soft |  | Scruffy |
| 2 | 5-10 | Loose stools/semiformed | Presence/hemoccult positive | Hunched and scruffy |
| 3 | 10-15 | Mild diarrhea (watery) |  | Non-motile, hunched, and scruffy |
| 4 | 15 or more | Gross diarrhea | Gross bleeding |  |

Consistent with previous studies [2,10,39], 2.5% DSS treatment in drinking water resulted in increases in DAI scores in both males and females in a time-dependent manner, with higher DAI scores as the days of treatment progressed. As expected, DAI scores in DSS-treated male and female mice were significantly (p < 0.0001) higher than scores in their respective water controls, validating that DSS treatment induces a symptom profile consistent with colitis in both sexes (**Figure 3C**). Notably, DSS-treated male mice displayed significantly (p = 0.0031) higher DAI scores when compared to female mice, demonstrating that male mice have worse clinical progression of colitis than female mice.

To dissect out the components within the cumulative DAI scores that contribute to the observed sex differences in the progression of colitis, we analyzed and compared each disease parameter individually in both male and female mice (**Figure 3D-G**). As predicted, DSS-treated male and female mice showed higher scores than their respective water controls in all the individual DAI components evaluated. Similar to the composite DAI scores, the progression of the individual disease components was also time-dependent, with higher scores observed with treatment progression in both males and females. While DSS-induced changes in body weight, physical appearance, and stool consistency were observed within a few days of treatment (**Figure 3D, E and F**), the presence of blood in the stool was not observed until days 4-6 after treatment (**Figure 3G**).

Comparison of the individual disease components in males and females further revealed that the sex differences observed in the composite DAI scores in DSS-treated mice arise from differences in the percentage of body weight loss and physical appearance scores but not from differences in the scores for stool consistency or fecal blood. Thus, as shown in **Figure 3D**, while both DSS-treated male and female mice lost weight during the days of treatment, the percentage of weight loss over time was significantly (p = 0.0193) higher in male than in female mice. Male mice also displayed worse DSS-induced changes in physical appearance than female mice, with a significantly (p = 0.0002) higher appearance score measured in male than in female mice (**Figure 3E**). In contrast, stool consistency and fecal blood scores were comparable in male and female mice for the duration of the DSS treatment (**Figure 3F-G**), demonstrating that these two disease components are not sex-dependent.

Evaluation of the time-course for DSS-induced disease progression further revealed that the onset of symptoms associated with colitis is also sex-dependent (**Table 2**). DSS-treated males, for example, exhibited significant (p = 0.0042) changes in cumulative DAI scores at treatment day 1 whereas female mice did not exhibit significant (p = 0.0157) changes until treatment day 2. Sex differences in symptom onset consistent with colitis were most pronounced in DSS-induced changes in body weight where males started to lose significant (p = 0.0124) weight from treatment day 1 while females did not display significant (p = 0.0098) DSS-induced weight loss until treatment day 7. Similarly, males start to exhibit significantly (p < 0.0001) worse appearance scores on day 2 of DSS treatment while the onset for significant (p = 0.0165) changes in appearance scores in females is on day 4. Consistent with the lack of sex differences in DSS-induced changes in stool consistency and presence of blood in the stool, analysis of the onset of changes in these two parameters was comparable in both sexes. Thus, significant (p < 0.05) DSS-induced changes in stool consistency scores were first observed on treatment day 2 and significant (p < 0.05) changes in fecal blood scores were observed on treatment day 6 in both male and female mice.

**Table 2.**
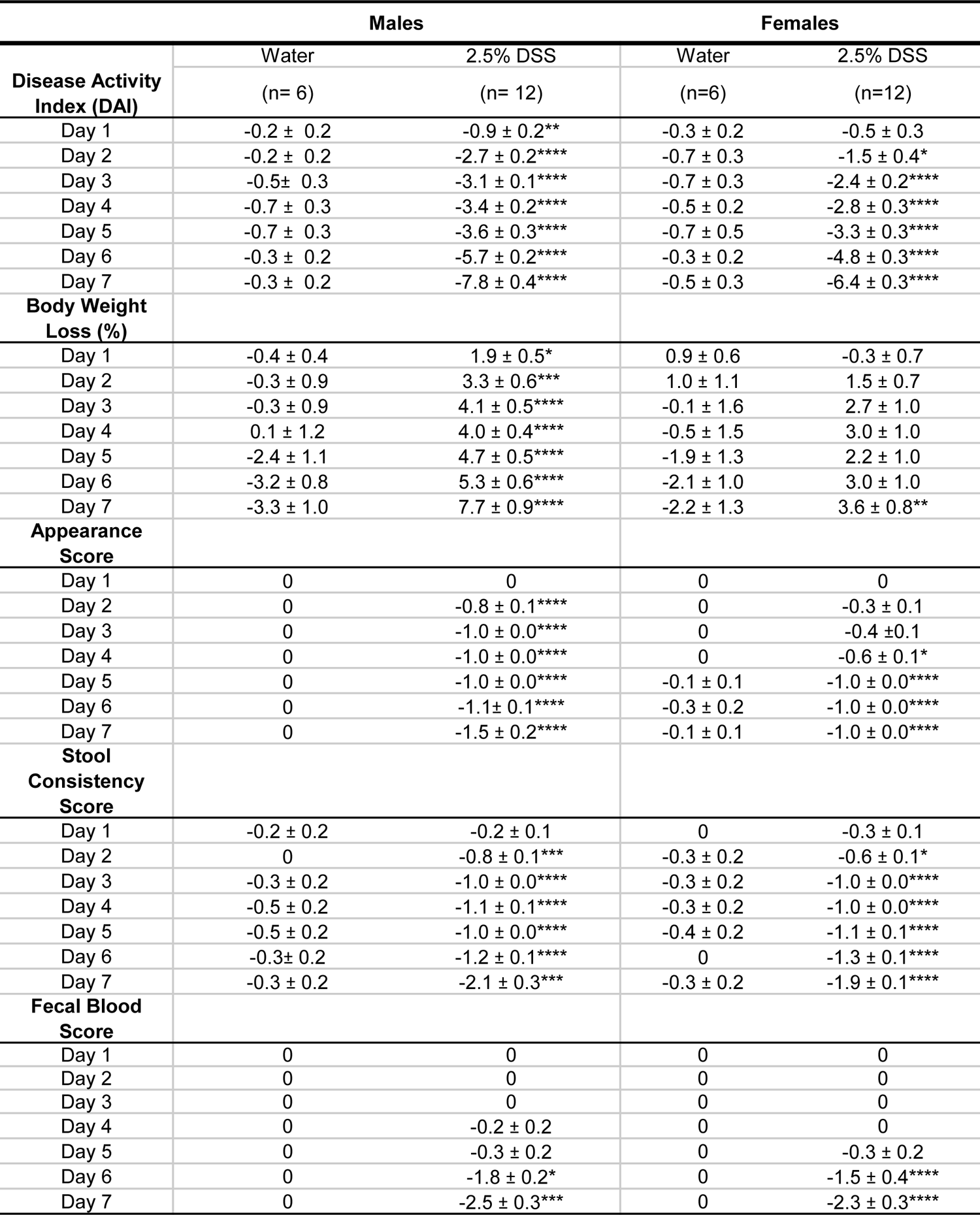
Onset of individual disease activity index (DAI) components following DSS treatment in male and female mice. Values are presented as the mean ± SEM difference from day 0 within the same treatment. *p≤0.05, **p≤0.01, ***p ≤ 0.001, ****p≤0.0001; Repeated Measures Two-Way ANOVA followed by Šídák’s multiple comparisons test compared to day 0.

Altogether, these results demonstrate that while both male and female mice treated with 2.5% DSS develop a clinical profile consistent with colitis, marked sex differences are observed in both the severity and onset of symptoms with males consistently exhibiting worse presentation of symptoms compared to females.

### Motivation to groom is comparable in both sexes after DSS-induced colitis

The experiments described above revealed worse physical appearance in DSS-treated males compared to females (**Figure 3E**). To assess if differential motivation in self-grooming behavior contributes to sex differences in DSS-induced changes in physical appearance, we performed the splash test in control and DSS-treated male and female mice while simultaneously scoring coat states to monitor changes in physical appearance as a function of time (**Figure 4A**). Immediately after sucrose solution application, all animals had a maximum coat state score independently of sex or DSS/water treatment (**Figure 4B**). The coat state in control male and female mice returned to baseline 50 minutes after sucrose solution application. Consistent with the appearance results shown in **Figure 3E**, the coat state of DSS-treated male and female mice did not return to baseline, displaying a scruffy and humid physical appearance for the duration of the experiment. Analysis of the total time spent grooming revealed, however, that control and DSS-treated male and female mice spend comparable time grooming (**Figure 4C**), suggesting that self-grooming motivation does not contribute to worse physical appearance in DSS-treated animals independently of sex.

**Figure 4.**
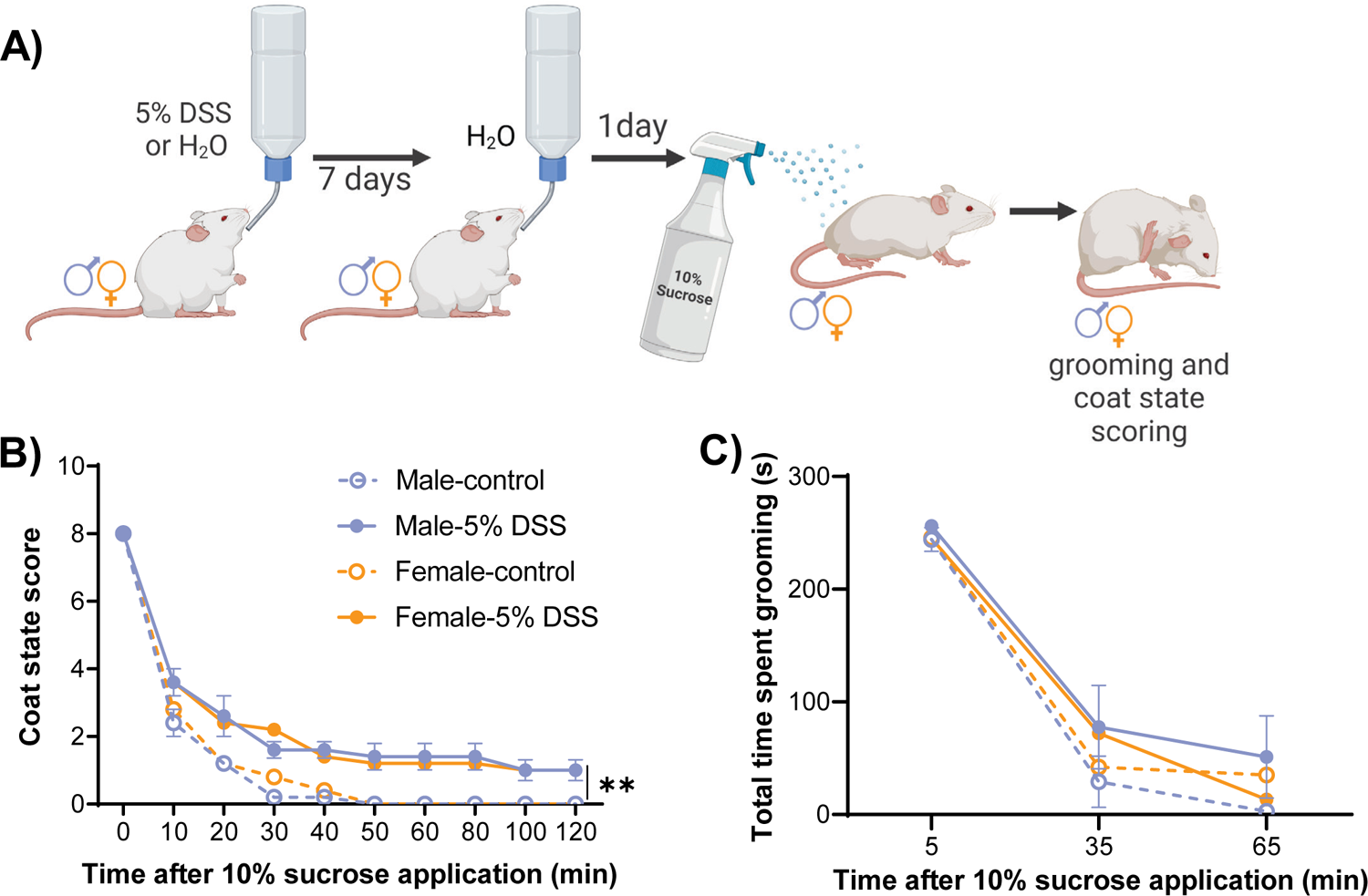
Motivation to groom is comparable in both sexes after DSS-induced colitis. (A) Timeline for splash test experiment. 10% sucrose solution was applied to the dorsal coat of control and DSS-treated male and female mice. (B-C) Coat state score (B), and total time spent grooming (C) were recorded. Data is presented as mean ± SEM. n = 5 females and 5 males. **p < 0.01: control vs 5% DSS; two-way ANOVA.

### Capsaicin-induced hypersensitivity is comparable in male and female mice after DSS-induced colitis, but the behavioral manifestation is sex-dependent

Altered pain perception in patients with Irritable Bowel Syndrome (IBS) and other functional gastrointestinal disorders has been associated with the development of visceral pain hypersensitivity [9, 36]. Thus, the model of intracolonic capsaicin was used to evaluate spontaneous visceral pain-related behaviors in male and female mice treated with 2.5% DSS as a model of IBS, IBD and other gastrointestinal disorders (**Figure 5A**). Consistent with the results observed in naïve female mice (**Figure 2**), DSS-treated females displayed a significant (p < 0.0001) increase in the total number of spontaneous nociceptive behaviors upon intracolonic administration of 0.01% capsaicin compared to respective control mice injected with vehicle (**Figure 5B**). Interestingly, unlike naïve male mice (**Figure 2**), DSS-treated males showed a significant (p < 0.001) increase in the total number of spontaneous nociceptive behaviors upon intracolonic administration of 0.01% capsaicin compared to those injected with vehicle control (**Figure 5B**).

**Figure 5.**
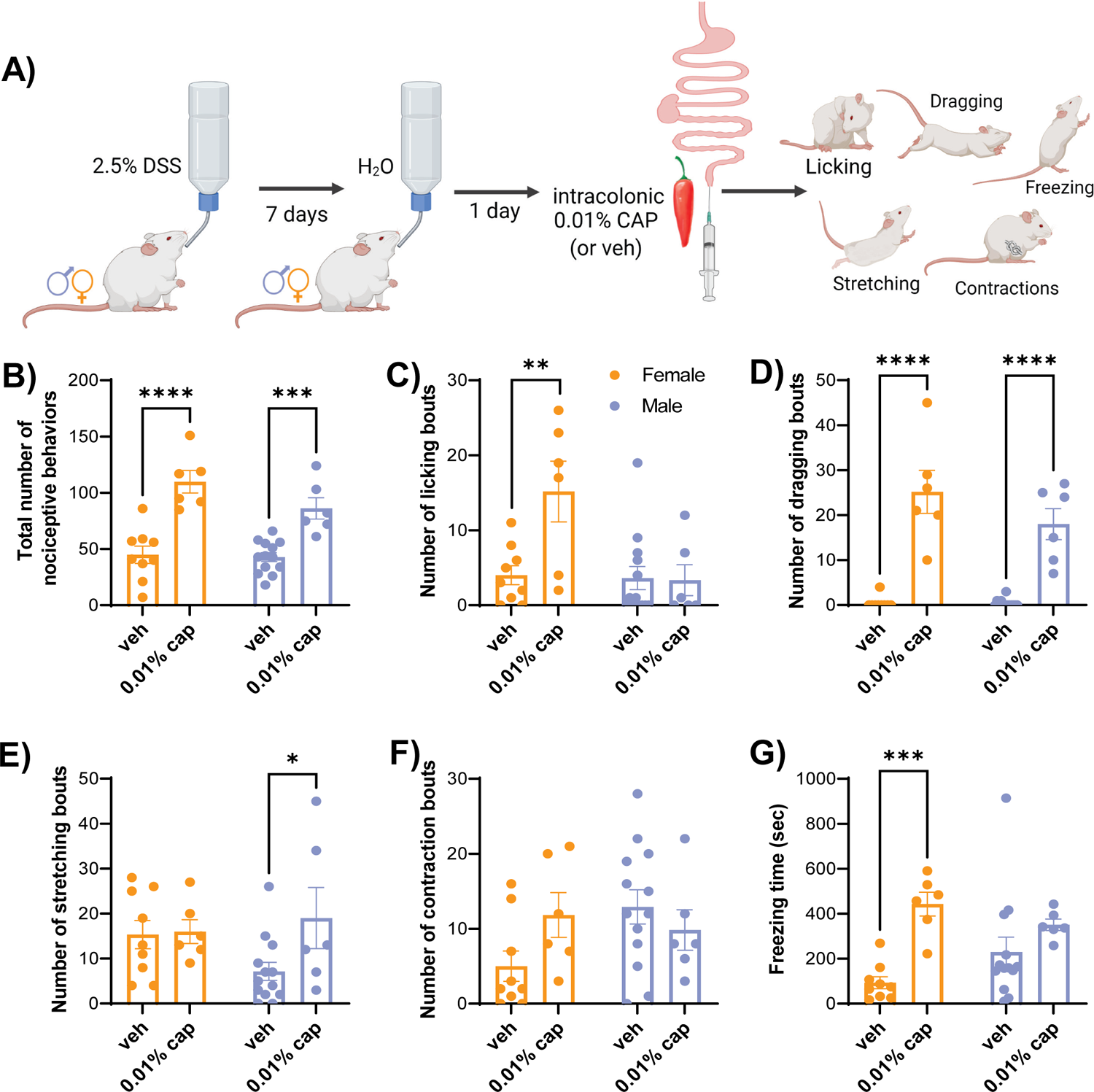
Capsaicin-induced hypersensitivity is comparable in male and female mice after DSS-induced colitis, but the behavioral manifestation is sex-dependent. (A) Timeline of behavioral experiments. DSS-treated male and female mice were injected with intracolonic 0.01% capsaicin and capsaicin-evoked nociceptive responses, defined as licking, stretching, dragging, and contractions of the abdomen, and freezing behaviors were recorded. Total number of pain-related behaviors (B), licking bouts (C), dragging bouts (D), stretching bouts (E), contraction bouts (F) and time spent in freezing behavior (G) in vehicle control and 0.01% capsaicin. Data is presented as mean ± SEM. n = 6-9 females and 6-13 males. *p < 0.05, **p < 0.01, ***p < 0.001, ****p < 0.0001: veh vs 0.01% cap; two-way ANOVA followed by Šídák’s multiple comparisons test.

To identify the specific pain-related behaviors contributing to capsaicin-induced visceral hypersensitivity observed in both sexes, the individual components that comprise the total number of pain-related behaviors were analyzed (**Figure 5C-F**). Consistent with the results observed in naïve female mice (**Figure 2**), DSS-treated females showed a significant (p < 0.01) increase in the number of capsaicin-induced licking (**Figure 5C**), dragging (**Figure 5D**) and freezing responses (**Figure 5G**) compared to vehicle-treated females while the number of contraction bouts (**Figure 5F**) were indistinguishable between capsaicin- and vehicle-treated females. In contrast, unlike the capsaicin-induced increases in stretching bouts seen in naïve female mice (**Figure 2**), capsaicin and vehicle-injected females displayed comparable stretching bouts after DSS treatment (**Figure 5E**). Our experiments in naïve male mice showed that both the cumulative and individual spontaneous pain-related behaviors where comparable in mice injected with 0.01% capsaicin or vehicle control (**Figure 2**). In contrast with these results, evaluation of the individual capsaicin-induced pain-related behaviors after DSS-induced colitis showed significant (p < 0.05) increases in the number of dragging (**Figure 5D**) and stretching bouts (**Figure 5E**) after intracolonic injection of 0.01% capsaicin, compared to intracolonic vehicle control injections. Consistent with the results observed in naïve male mice, however, the number of licking (**Figure 5C**) and contraction bouts (**Figure 5F**) as well as time spent freezing (**Figure 5G**) were indistinguishable in capsaicin and vehicle-injected DSS-treated males. Collectively, these results demonstrate that both sexes develop capsaicin-induced hypersensitivity to intracolonic application of 0.01% capsaicin after DSS-induced colitis. The manifestation of hypersensitivity, however, was sex-dependent, with both sexes displaying increases in dragging behavior (**Figure 5D**) but only females showing increases in licking and freezing behaviors (**Figure 5C and 5G**) and only males displaying increases in stretching behaviors (**Figure 5E**).

### Referred abdominal hypersensitivity is sex-dependent after acute but not persistent colon inflammation

Previous studies have shown that following intracolonic administration of capsaicin or DSS-induced colitis, mice display referred abdominal hypersensitivity, manifested as an increase in the frequency of pain-related responses to tactile stimulation of the abdomen during the von Frey Test (Laird, Martinez-Caro et al. 2001, Jain, Hassan et al. 2015). The next set of experiments aimed at evaluating potential sex differences in baseline responses to tactile stimulation of the abdomen using a 0.16g von Frey filament as well as in referred abdominal hypersensitivity after acute and persistent colon inflammation, induced by intracolonic administration of 0.01% capsaicin or DSS in drinking water, respectively (**Figure 6A**). Response frequency was indistinguishable between males and females in control conditions (**Figure 6B**), demonstrating that baseline responses to tactile stimulation of the abdomen are not sex dependent. Consistent with previous reports [29], intracolonic injection of 0.01 % capsaicin significantly (p < 0.001) increased response frequency to tactile stimulation of the abdomen in female mice compared to intracolonic vehicle control injections (**Figure 6B**). In contrast, response frequencies were indistinguishable in male mice following intracolonic administration of 0.01% capsaicin or vehicle control. These findings are consistent with the higher capsaicin-induced spontaneous nociceptive visceral responses we observe in females (**Figure 2**), further demonstrating that referred abdominal hypersensitivity is also sex-dependent in the context of acute colon inflammation.

**Figure 6.**
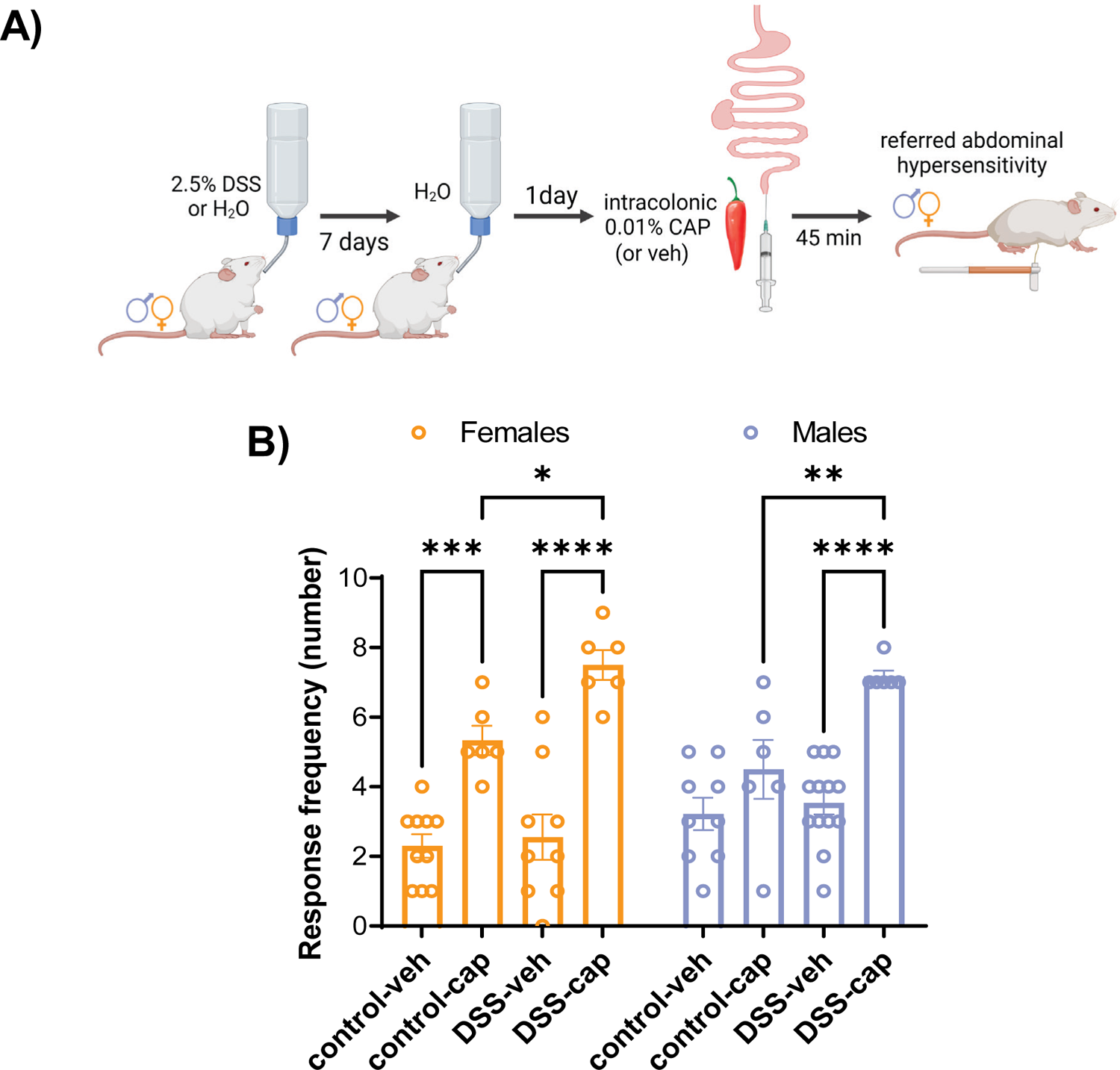
Referred abdominal hypersensitivity is sex dependent after acute but not persistent colon inflammation. (A) Timeline for von Frey behavioral test in the context of acute and persistent colon irritation. Control and DSS-treated male and female mice were injected with intracolonic 0.01% capsaicin or vehicle control and abdominal sensitivity to tactile stimulation was measured. (B) Quantification of response frequency to tactile stimulation of the abdomen in control and DSS-treated male and female mice after intracolonic injection of 0.01% capsaicin or vehicle control. Data is presented as mean ± SEM. n = 6-13 males and 6-10 females. *p < 0.05: control-0.01% cap vs DSS-0.01% cap, **p < 0.01: control-0.01% cap vs DSS-0.01% cap, ***p < 0.001: control-veh vs control-0.01% cap, and ****p < 0.0001: DSS-veh vs DSS-0.01% cap; two-way ANOVA followed by Tukey’s multiple comparisons test.

Evaluation of DSS-induced referred abdominal hypersensitivity in males and females revealed that both male and female mice displayed significant (p < 0.0001) increases in response frequencies to tactile stimulation of the abdomen using a 0.16g von Frey filament after intracolonic administration of 0.01% capsaicin when compared to animals injected with intracolonic vehicle control (**Figure 6B**). Further analyses showed that, as expected, capsaicin-induced hypersensitivity to tactile stimulation is potentiated by DSS-induced persistent colon inflammation, with both sexes showing a significant (p < 0.05) increase in response frequencies to tactile stimulation of the abdomen after intracolonic administration of 0.01% capsaicin in the context of DSS-induced colitis when compared to control mice injected with 0.01% capsaicin. These combined results are consistent with our findings showing that spontaneous capsaicin-induced visceral responses in the context of DSS-induced colitis is not sex-dependent (**Figure 5**) and further demonstrate that referred abdominal hypersensitivity in the context of persistent colon inflammation is also not sex-dependent.

### DSS-induced bowel pathology is comparable in males and females

The results presented above show that clinical progression of disease and behavioral manifestation of hypersensitivity following DSS treatment are sexually dimorphic. To gain insight into whether DSS-induced bowel pathology contributes to the observed sex differences, colon lengths and histopathological analysis of the bowels of control and DSS-treated mice of both sexes were evaluated (**Figure 7A**). Bowel length is commonly used to evaluate macroscopic manifestations of gastrointestinal disease in rodents, with shortening of the bowel reported in many gastrointestinal conditions, including DSS-induced colitis [2, 39]. Consistent with previous reports, analysis of our experiments showed that both sexes exhibit significant (p < 0.0001) shortening of the bowel after DSS treatment when compared to their respective water controls (**Figure 7B-C**). Further analysis revealed that female bowels are significantly (p < 0.05) shorter than male bowels in both control and DSS-treated mice. Comparison of the change in average bowel length in control and DSS-treated mice reveals, however, that both sexes display a bowel shortening of ∼19% after DSS treatment, when compared to their respective water controls (**Figure 7C**). These results demonstrate that the manifestation of DSS-induced colitis is comparable in males and female mice at the gross pathological level.

**Figure 7.**
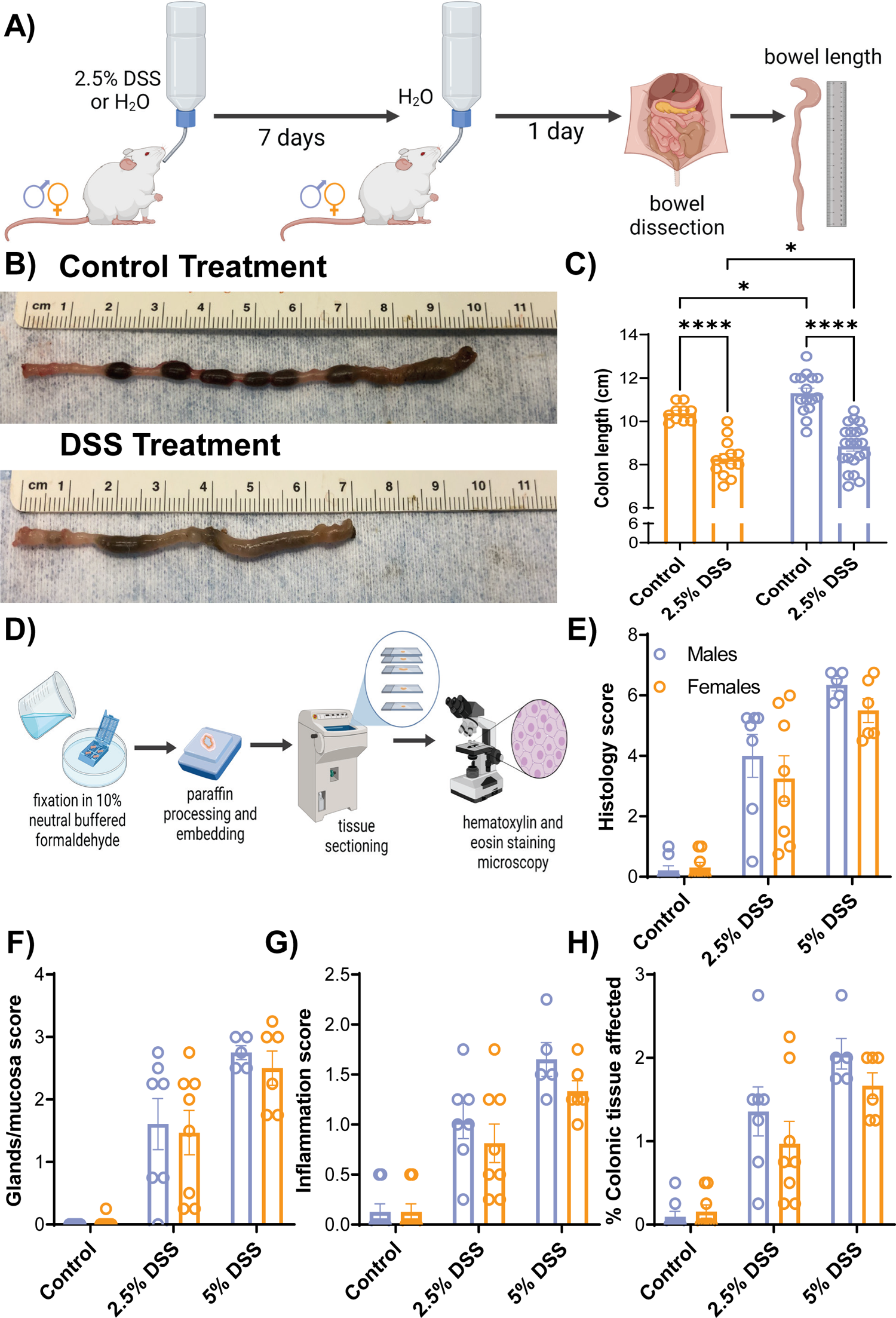
DSS-induced bowel pathology is comparable in males and females. (A) Timeline for bowel pathology analysis. Bowels of control and DSS-treated male and female mice were dissected, colon length was measured 1d after the end of DSS treatment and colonic samples were collected for histopathological analysis. (B) Representative images of control (top) and DSS-treated (bottom) colon samples. (C) Colon length in control and DSS-treated male and female mice. (D) Timeline for histological analysis of DSS-induced colon pathology. (E) Cumulative histology score in bowels from control and DSS-treated male and female mice and its individual components, defined as glands/mucosa score (F), inflammation score (G) and % colonic tissue affected (H). Data is presented as mean ± SEM. n = 6-13 females and 5-21 males. *p<0.05: males vs females and ****p<0.0001: control vs 2.5% DSS; two-way ANOVA followed by Tukey’s multiple comparisons test.

Histopathological analysis of the colon of both control and DSS-treated male and female mice was performed to assess bowel pathology at the microscopic level (**Figure 7D-H**). Analyses of the cumulative histology score, composed of glands-mucosa, inflammation and % of colonic tissue affected scores, revealed that DSS induces increases in histology scores in bowels from both male and female mice in a dose-dependent manner, with higher histology scores associated with increasing dose of DSS administered (**Figure 7E**). Consistent with our analysis of gross bowel pathology, analyses of histology scores showed comparable pathology at the microscopic level between DSS-treated male and female mice. Analyses of the individual components further demonstrated comparable damage of the colonic crypts and mucosa (**Figure 7F**), inflammatory infiltration of leucocytes (**Figure 7G**) and % of colonic tissue affected (**Figure 7H**) in DSS-treated males and females. Altogether, these results show no measurable sex difference in DSS-induced gross and microscopic colon pathology.

## DISCUSSION

Chronic visceral pain is predominantly reported in women and higher pain sensitivity has been shown in women patients [46]. Despite this, the majority of preclinical pain studies have focused only on males [35]. The limited inclusion of sex as a biological variable in preclinical studies results in an incomplete understanding of the biological processes underlying pain [35]. In this study, we evaluated and characterized potential sex differences in visceral pain-related responses, disease progression of colitis and colitis-induced colon pathology using the intracolonic capsaicin and DSS-induced colitis models of chemically induced visceral hypersensitivity in mice (**Figure 8**). Our behavioral experiments show sex-dependent differences in visceral pain-related behaviors, with females exhibiting more pain-related responses and referred abdominal hypersensitivity than males in the context of acute colon inflammation induced by intracolonic capsaicin (**Figure 8A-B**). We further show that colitis-induced visceral hypersensitivity and referred abdominal hypersensitivity are similar between males and females, but the manifestation of capsaicin-induced behavioral responses is sex-dependent. While both sexes exhibited dragging, stretching, and freezing in response to intracolonic capsaicin, increases in licking of the abdomen were only observed in females and increases in abdominal contractions were only displayed by males (**Figure 8A**). Consistent with other studies [2,10,39], we also show that progression of DSS-induced colitis is sex-dependent, with males exhibiting worse clinical progression, manifested as worse appearance and higher weight loss, than in females (**Figure 8C**). No measurable sex differences were observed, however, in colitis-induced bowel pathology, stool consistency or fecal blood (**Figure 8D**). Together, these findings stress the importance of incorporating both sexes when studying mechanisms that drive clinical and behavioral manifestations of visceral hypersensitivity.

**Figure 8.**
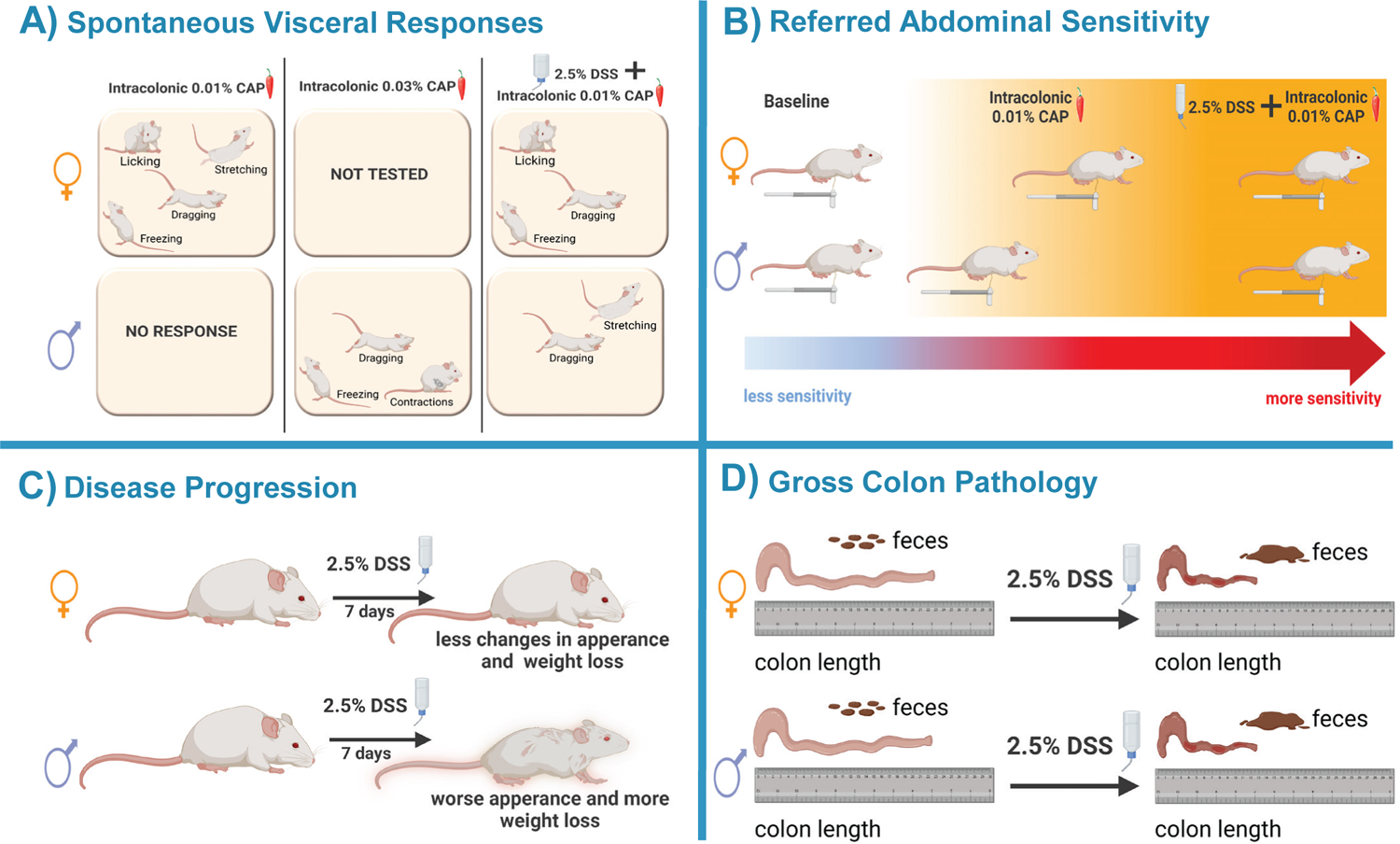
Summary of main findings. (A) Behavioral manifestation of capsaicin-induced spontaneous visceral nociceptive responses is sex dependent after acute and persistent colon inflammation. After intracolonic capsaicin, females exhibit more capsaicin-induced nociceptive behaviors than males. In the context of DSS-induced colitis, visceral hypersensitivity is comparable in both sexes, but the manifestation is distinct. (B) Referred abdominal hypersensitivity is higher in females after intracolonic capsaicin but is similar between sexes after persistent colon inflammation or at baseline. (C) Progression of DSS-induced colitis is sex dependent. Worse physical appearance and greater weight loss is observed in males compared to females. (D) No measurable sex difference was observed in DSS-induced bowel shortening.

### Behavioral manifestation of visceral pain-related behaviors is sex-dependent

While previous studies consistently show that the model of DSS-induced colitis and intracolonic capsaicin produce visceral hypersensitivity and referred abdominal hyperalgesia in both sexes [20, 29], potential sex differences in pain-related behaviors in these two models of visceral pain have not been evaluated. In the present study, we demonstrated that visceral pain-related behaviors in freely behaving mice is sex-dependent. We specifically showed that females display higher capsaicin-induced spontaneous nociceptive responses than males and that the behavioral manifestation of pain-related responses to intracolonic capsaicin is also dependent on sex, with increases in licking of the abdomen observed only in females and increases in abdominal contractions only displayed by males (**Figures 2 and 5**).

Sex differences in behavioral outputs have been described previously in other contexts such as fear and escape behaviors [12,14,27], highlighting the importance of characterizing behavioral assays in both males and females. Previous studies have further shown that neural circuits driving distinct pain-related responses, such as paw withdrawal or licking in response to evoked tactile stimulation, are unique [17]. Our behavioral experiments showing that male and female mice exhibit distinct behavioral responses to intracolonic capsaicin suggest that recruitment of neural circuits in responses to a particular noxious stimulus might be sex-dependent.

### Clinical progression of disease in a mouse model of colitis are sex-dependent

Our experiments showed that DSS treatment induces a clinical profile consistent with colitis in both male and female mice and that males exhibit a worse clinical progression of disease (**Figure 3**). The disease activity index (DAI) used to measure clinical progression of colitis is typically presented as a cumulative score of the following components: appearance, weight loss, stool consistency and fecal blood [2,6,10,25,32,39]. Analysis of the individual DAI components in the present study revealed that higher DAI scores in males are driven by greater weight loss and worse appearance but that DSS-induced changes in stool consistency and fecal blood are similar between sexes. The lack of measurable sex differences in stool consistency and fecal blood, together with the comparable DSS-induced shortening of the bowel (**Figure 7**) suggests that gross pathology of the bowel is not the driving force behind the sex differences in weight loss. Weight loss is a hallmark in animal models of experimental colitis and has been associated with decreases in food intake and/or changes in metabolic rates, locomotor activity or body fat content [33]. Similarly, weight loss in IBD patients has been related with decreased appetite, alterations in metabolic hormones and/or malabsorption of nutrients [7]. Future studies to explore whether DSS induces sex-dependent changes in food consumption and/or metabolic alterations that contribute to colitis-induced weight loss are needed.

Despite the worse physical appearance seen in DSS-treated males (**Figure 3E**), coat state and time spent grooming after sucrose solution application in the splash test was similar between males and females (**Figure 4**). These results are consistent with findings from previous studies in males [20] and suggest that motivation to self-groom is not affected after DSS-induced colitis and it is not sex-dependent. When interpreting these findings, it is important to consider that sucrose solution application creates an acute challenge that experimentally evokes grooming. Motivation to groom when presented with an acute challenge like sucrose application might be different to motivation to self-groom for daily upkeeping. Previous studies have shown that self-grooming is an evolutionary conserved and innate behavior in mice that involves a complex sequencing pattern and contributes to hygiene maintenance, thermoregulation, social communication, and other biological processes [24]. The worse physical appearance in DSS-treated males, combined with the lack of sex or treatment-dependent effect on time spent grooming in response to sucrose application, suggest that males are less efficient at grooming than females after the induction of colitis.

An important caveat of the experiments presented here is that the same percentage of DSS was used in males and females, despite males having greater initial body weight than females. Consequently, the dose of DSS per body weight is most likely lower in males than it is in females. Whether sex differences in metabolism and excretion rates of DSS influence the observed DSS-induced phenotypes is also unknown. Despite potential differences in the absolute DSS dose intake, however, males exhibit worse disease progression than females, further emphasizing sexual dimorphism in disease progression in the context of colitis.

### Comparison with other studies and potential translational application

Our behavioral experiments show that female mice display more capsaicin-induced spontaneous visceral pain-related responses than males (**Figure 2**). These results are consistent with previous studies showing that female rats have higher visceromotor responses to noxious colorectal distension at baseline and after intracolonic injection of mustard oil than male rats [15,22,23]. Measurements of visceromotor responses to colorectal distension are typically performed in lightly anesthetized or loosely restrained rats, which may confound the behavioral outcomes measured and limit the interpretation of findings. The consistencies between our behavioral findings in freely behaving mice and those in previous studies using lightly anesthetized or loosely restrained rats demonstrate that sex differences in visceral pain-related behaviors are comparable and can be equally studied in freely behaving mice or in rats under light anesthesia or loosely restrained. Our results showing that female rodents display higher visceral sensitivity than males are also consistent with clinical studies in humans that also show higher visceral sensitivity in women than men [3,4,44], validating the use of rodent preclinical models to mechanistically study sex differences in visceral pain. Lastly, the demonstration that sex differences in visceral pain-related responses are recapitulated in mice is important as they offer a more enriched repertoire of molecular genetic tools to study behavior at circuit and mechanistic levels.

It is important to note that our experiments showed that visceral hypersensitivity in the context of DSS-induced colitis is not sex-dependent (**Figure 5**). These findings contrast with clinical studies that report that female IBS patients exhibit higher visceral sensitivity and report more severe abdominal pain or discomfort than men (Chang, Mayer et al. 2006, Tang, Yang et al. 2012, Camilleri 2020). As discussed above, however, our experiments also demonstrated that the behavioral manifestation of visceral pain-related responses in mice is sex-dependent (**Figure 2 and 5**), suggesting that pain perception is differentially experienced by females and males. Given that pain is a multidimensional experience that comprises affective, cognitive, and somatosensory components, studies that evaluate these components in humans are needed to understand the mechanisms driving sex differences in visceral pain. Based on these results, improved diagnostic and therapeutic options for pain relief can be developed.

At a pathological level, consumption of DSS in drinking water has been shown to disturb the colonic lumen structure, eliciting an inflammatory response that emulates inflammatory bowel diseases (IBD) such as ulcerative colitis (UC) and Crohn’s disease (CD) [30]. Consistent with previous reports [39], we also show that DSS induces shortening and histopathological changes in the colon (**Figure 7**) that resemble the shortening of the small bowel and histopathological features observed in patients with ulcerative colitis (UC) and Crohn’s disease (CD) [38], further validating the applicability of this model to study IBDs. Notably, however, while previous studies have reported sex differences in DSS-induced bowel shortening and histopathology [2,10,31], our experiments showed that bowel pathology is comparable between sexes. These discrepancies could be due to differences in strain used, DSS supplier or animal facility-specific factors that have been previously shown to strongly influence DSS experiments [5, 31].

The experiments in the present study provide evidence for sex differences in visceral pain-related behaviors and clinical progression of colitis in preclinical models of acute and persistent colonic irritation. Together, these results highlight the importance of considering sex as a biological variable in both preclinical and clinical pain studies.

## MATERIALS AND METHODS

### Subjects

All animal procedures were performed in accordance with the guidelines of the National Institutes of Health (NIH) and were approved by the Animal Care and Use Committee of the National Institute of Neurological Disorders and Stroke and the National Institute on Deafness and other Communication Disorders. Adult male and female Swiss-Webster mice (Taconic Farms) between 8-13 weeks old were used for all experiments. Littermates were randomly assigned to experimental groups and group housed (up to 5 mice per cage) under reversed 12 h light/dark cycle (9 pm to 9 am light). One week prior to experiments, mice were housed in pairs in clean home cages separated by a perforated Plexiglas divider. Food and water were provided ad libitum. Prior to all behavioral experiments, mice were handled as previously described for at least 5 days to minimize potential stress effects associated with handling [18]. Male and female mice were never tested simultaneously in the same behavior room. All experimental procedures were performed by an observer blind to experimental treatment and replicated at least 3 times.

### DSS-induced Model of Colitis

Male and female mice were randomly assigned to drinking water (control group) or 2.5% (wt/vol) Dextran Sulfate Sodium (DSS) salt (reagent-grade, mol. wt. 36 to 50 kDa; MP Biomedicals, #16011050) dissolved in drinking water to induce colitis (experimental group), ad libitum for seven days. On day 7, mice were switched to regular water ad libitum for the duration of the experiment. Intracolonic capsaicin, referred abdominal sensitivity and bowel sample collection experiments were performed on day 8 (**Figure 1A**). Each day, along with the handling of the mouse, the disease activity index (DAI) was logged by an investigator blind to experimental treatment to evaluate and score disease progression as previously described (Kim et al., 2012). The DAI is a composite of scores for weight loss, appearance, stool consistency, and blood presence in the stool using a Hemoccult card (Beckman Coulter). See **Table 1**.

### Intracolonic Capsaicin

On experiment day 8 (see full experimental timeline above and in **Figure 1A**), male or female mice were habituated (for 1hr) and tested on an elevated mesh platform in individual 11 × 11 × 13 cm ventilated opaque white Plexiglas boxes. A mirror was placed at 45-degree angle under the mesh platform to allow full visualization of the animals. Intracolonic injections were performed as previously described [40]. Briefly, mice were anesthetized with 1% isoflurane in an induction chamber, and then kept lightly anesthetized with 0.5%–1% isoflurane at a flow rate of 0.5 L/min. A light layer of petroleum jelly (Vaseline) was applied to the peri-anal area and tubing to avoid the stimulation of somatic areas and ease tube insertion. Capsaicin (0.01% or 0.03%) was prepared using a stock solution of capsaicin (Sigma Aldrich) diluted in ethanol, Tween80, and saline (10/10/80, respectively). 50µl of capsaicin (0.01% or 0.03%) or vehicle control (10% ethanol, 10% Tween 80, and 80% saline) was slowly injected via PE-10 non-toxic, sterile polyethylene tubing (0.28 mm ID / 0.61 mm OD; Daigger Scientific) with a rounded tip connected to a blunted 30G x 1/2 needle (BD PrecisionGlide) and 1 cc syringe (Terumo) that was gently introduced 4 cm into the colon via the anus. Spontaneous nociceptive responses to intracolonic capsaicin (or vehicle control) were measured for 20 minutes immediately following intracolonic injections. Spontaneous nociceptive responses were defined as licking of abdomen, stretching of abdomen, dragging and abdominal contractions. The time spent freezing following the injection was also measured during the 20-minute test period.

### Referred Abdominal Hypersensitivity

Referred abdominal hypersensitivity was evaluated 45 minutes after capsaicin (or vehicle control) intracolonic injection as described previously [40]. The abdominal hair of all test mice was removed (24 h) before testing with a depilatory (Nair). A 0.16g von Frey filament (North Coast Medical, Inc. San Jose, CA) was applied to the abdomen for approximately 1–2 s, with a stimulus interval of 15 s. Positive responses were defined as a rapid withdrawal, jumping, licking or abdominal contractions immediately following application of the von Frey filament to the abdomen. A total of 10 trials were performed and the number of positive responses per animal were recorded and reported as response frequency.

### Splash Test

On experiment day 8 (see full experimental timeline above and in **Figure 1A**), control and DSS-treated male and female mice were individually transferred to a new home cage with regular bedding and were habituated in them for at least 1hr. Each mouse received two sprays of 10% sucrose solution on their dorsal coat. Immediately following sucrose solution application, the coat state of the animals was scored every 10 min for 1hr and every 20 min starting at the 1-hour timepoint until the 2-hours timepoint from the sucrose solution application. The coat state score was obtained using the following system: 8 = wet, soaked dorsal coat; 4 = scruffy and humid dorsal coat; 2 = dry and smooth lower back coat but upper back coat is still scruffy and/or humid; 1 = mostly dry and smooth dorsal coat with a few scruffy patches and 0 = completely dry and smooth dorsal coat. The time spent in grooming behavior was measured during a five-minute period immediately after sucrose solution application as well as 35 and 65 minutes after sucrose solution application. The observer was always blind to experimental treatment.

### Bowel Sample Collection and Histopathological Analysis

On experiment day 8 (see full experimental timeline above and in **Figure 1A**), control and DSS-treated males and females were anesthetized with 1% isoflurane in an induction chamber and euthanized by cervical dislocation. Bowels were dissected and the colon was carefully removed to measure its length. The colon was subsequently divided in proximal, medial, medial/distal, and distal 1 cm sections, fixed in 10% Neutral Buffered Formaldehyde (Azer Scientific) for 24hrs and stored in 70% ethanol until processing. Tissue samples were embedded in paraffin blocks, cut in 5µm sections, fixed to glass slides, and stained with hematoxylin and eosin (H&E) by Histoserv (Germantown, MD). Histopathological analysis of the tissue samples was performed as previously described [8, 25]. Tissue damage was assessed using a cumulative histology score of glands-mucosa, inflammation and % of colonic tissue affected. The score for glands-mucosa was as follows: 0 = no changes in glands-mucosa structure; 1 = loss of up to 1/3 of gland, crypts lifted off muscularis mucosae; 2 = loss of 2/3 of gland, crypts lifted off muscularis mucosae, little inflammation, thinning and loss of epithelial cells in the *intact* glands, cryptitis; 3 = loss of all the glands but the superficial epithelium is intact, mild infiltrate; 4 = erosion, ulceration, mild - moderate infiltrate. Inflammation score was defined as 0 = none to few leucocytes; 1 = mild, some increase in leucocytes at tips of crypts; 2 = moderate; 3 = severe, dense infiltrate throughout, transmural. Percent of colonic tissue affected was scored as 1 = 1 – 25%; 2 = 25 – 50%; 3 = 50 – 75%; 4 = 75 – 100 %. Analyses of the histology score and its individual components were initially performed separately per anatomical section of the colon collected (i.e. proximal, medial, medial/distal, and distal). This analysis showed that DSS-induced histopathology is comparable between all colon sections evaluated within bowels from mice of the same sex and dose of DSS administered. Based on these results, we calculated the average score of all 4 colonic sections per mouse for each parameter and used these values for subsequent graphs and analyses.

## Statistics

Results are expressed as mean ± standard errors of the mean (SEM). Analysis was performed using either Unpaired t-tests (without Welch’s correction for variance), Mann-Whitney U tests, two-way analyses of variance (ANOVA) or two-way analyses of variance (ANOVA) followed by Tukey’s multiple comparison tests or by Šidák’s multiple comparison. All analyses were performed using GraphPad Prism (v. 9) and p values lower than 0.05 were considered significant and are reported in figure legends.

## Data Availability

All data in this study is available from the corresponding author.

## AUTHOR CONTRIBUTIONS

Conceptualization, C.P., S.M.G., Y.C., A.M.F.M; Investigation, S.M.G., T.D.W., L.G.R., A.M.F.M; Data Analysis, A.M.F.M, S.M.G., T.D.W., L.G.R., L.R.B., Y.C.; Writing – Original Draft, A.M.F.M, Y.C.; Writing – Review & Editing,; Supervision, Y.C.; Funding Acquisition, Y.C.

## ACKNOWLEDGEMENTS

This research was supported by the National Center for Complementary and Integrative Health Intramural Research Program. We would like to thank Dr. David Shurtleff for feedback on this study, Uriah Contreras for technical assistance in the project and Dr. Mark Pitcher for training and assistance with the behavioral experiments. We would also like to thank the National Institutes of Neurological Disorder Mouse Facility staff for their vital work in animal husbandry. Cartoons in Figures 1-7A, 7D and 8 were created with BioRender.com.

